# Humans Use Forward Thinking to Exert Social Control

**DOI:** 10.1101/737353

**Authors:** Soojung Na, Dongil Chung, Andreas Hula, Jennifer Jung, Vincenzo G. Fiore, Peter Dayan, Xiaosi Gu

**Affiliations:** The Graduate School of Biomedical Sciences, Icahn School of Medicine at Mount Sinai; Nash Family Department of Neuroscience, Icahn School of Medicine at Mount Sinai; Department of Psychiatry, Icahn School of Medicine at Mount Sinai; Department of Human Factors Engineering, Ulsan National Institute of Science and Technology; Austrian Institute of Technology; School of Behavioral and Brain Sciences, The University of Texas at Dallas; Max Planck Institute for Biological Cybernetics, Tübingen, Germany; James J. Peter Veterans Affairs Medical Center, Bronx, NY

## Abstract

Social control, the ability to exert influence over others, is critical in interpersonal interactions yet uninvestigated. Here, we used functional neuroimaging and a social exchange paradigm in which people’s current choices either did, or did not, influence their partners’ proposals in the future. Computational modeling revealed that participants used forward thinking and calculated the downstream effects of their current actions regardless of the controllability of the social environment. Furthermore, greater levels of estimated control correlated with better performance in controllable interactions and less illusory beliefs about control in uncontrollable interactions. Neural instantiation of trial-by-trial values of social controllability were tracked in the ventromedial prefrontal cortex (vmPFC), striatum, and insula for controllable interactions, but only in vmPFC for uncontrollable interactions. These findings demonstrate that humans use forward thinking, a strategy similar to model-based planning, to guide social choices; and that subjective beliefs about social controllability might not be grounded in reality.

## INTRODUCTION

Power and control are central themes of almost all human interactions throughout history. Being in control is almost always preferred. One reason is that it can indicate one’s superior social status in a hierarchical structure such as being the king in a monarchy or the CEO of a company. However, even in non-hierarchical relationships, social controllability can confer actual benefits such as access to more resources. One example is business negotiations, where controllability is crucial for exerting influence over one’s counterparts to achieve desired monetary outcomes. Furthermore, controllability in general has been also associated with better mental health outcomes such as higher subjective well-being (Lachman and Weaver, 1998) and less negative affect(Maier and Seligman, 2016; Southwick and Southwick, 2018). However, the neurocomputational mechanisms that underlie how individuals exert control over others during social interactions have not been examined.

We hypothesize that social controllability is implemented through forward thinking in humans. This hypothesis is based on a major advance in recent decision-making literature – the emergence of computational and normative models demonstrating how people use ‘model-based’ (MB) planning to simulate the future and exert behavioral control, rather than merely relying on reinforcement learning of cached values alone (so-called ‘model-free’ planning) (Daw et al., 2011; Dolan and Dayan, 2013; Gläscher et al., 2010). Similar concepts have been applied to model iterative interactions such as the trust game (Hula et al., 2015), where people plan future interactions with the same partner. In social contexts, one’s current choices could contribute to one’s reputation and thus have a long-lasting effect on future social interactions. Thus, we predict that this mode of forward control would be crucial for strategic social interactions with multiple partners as well, such that a social agent takes into account not only decision variables related to the present, but also those related to the future.

In forward planning during social interaction, one’s subjective belief about the level of control could also have a key role. Given its importance, we might hope that individuals derive their beliefs about controllability from actual interactions. However, it is well established that human beliefs need not be perfectly calibrated or rooted in reality (Langer, 1975). Pertinent to controllability, previous studies have shown that overly optimistic beliefs about control in environments with little or no controllability (i.e., an illusion of control (Langer, 1975)) could impede performance in gamblers (Langer, 1975), traders (Fenton-O’Creevy et al., 2003), and drivers (Stephens and Ohtsuka, 2014). In contrast, pessimistic beliefs of no control when environments are controllable (i.e., hopelessness or helplessness) have also been associated with negative affect and relevant neural circuits (Maier and Seligman, 2016). Indeed, subjective beliefs about social controllability might not necessarily be grounded in the actual controllability of one’s social interactions; such a disconnection could also be counterproductive and even, in extreme cases, be indicative of psychiatric symptoms (Brown and Siegel, 1988; Harrow et al., 2009). Our second hypothesis is that during social interactions, humans’ subjective beliefs about control could also be an independent dimension from the actual controllability exerted in these interactions.

We used computational modeling, functional magnetic resonance imaging (fMRI), and a social exchange paradigm (see **Figure 1** and **Methods**), to test the two aforementioned hypotheses – 1) forward thinking serves as a mechanism for social control, and 2) actual social controllability and subjective beliefs about control (i.e. based on self-reports) could be two independent dimensions. Forty-eight healthy individuals participated in the task where they could (‘In Control’; 40 rounds) or could not (‘No Control’; 40 rounds) influence their partners’ proposals of monetary offers in the future (see **Figure 1a,b**, and **Methods** for details). Participants were told that they were playing members coming from two different teams, one each for the two control conditions (in a counterbalanced order across subjects); in fact, they played with a computer algorithm in both cases. **Notes S1** provides the task instruction provided to participants; **Figure S1a-c** describes a control experiment where participants were explicitly told they were playing against a computer algorithm.

**Figure 1.**
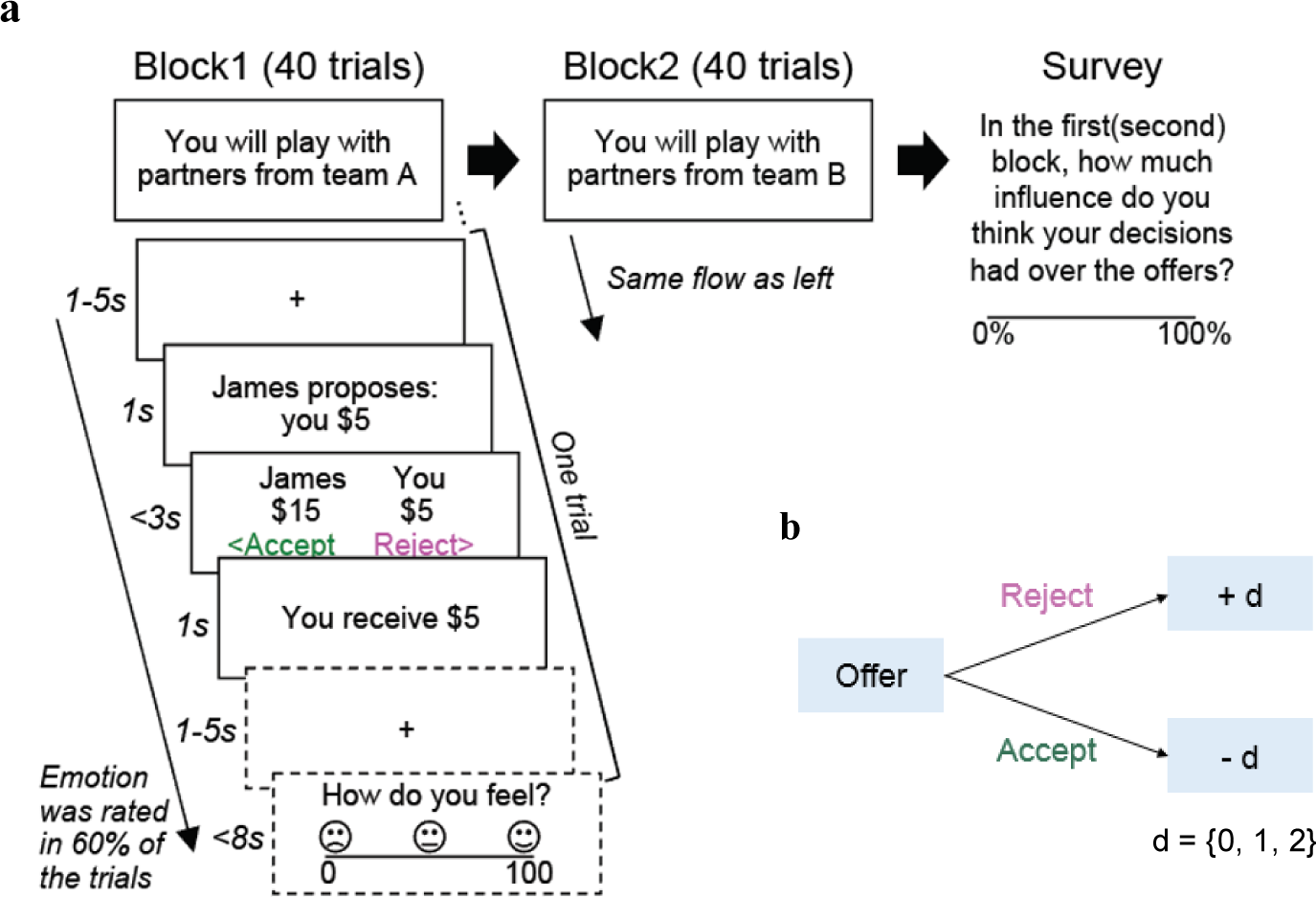
Experimental paradigm. (**a**) Participants played a social exchange task based on the ultimatum game. There were two blocks, one ‘In Control’ condition and the other ‘No Control’ condition. The order of the condition was counterbalanced. Each block had 40 trials. In each trial, participants needed to decide whether to accept or reject the split of $20 proposed by virtual members of a team. Participants rated their emotions after their choice in 60% of the trials. Upon the completion of the game, participants rated their subjective beliefs about control for each block. (**b**) The schematic of the offers (the proposed participants’ portion of the split) generation under In Control condition. Under the In Control condition, if participants accepted the offer at trial *t*, the next offer at trial *t+1* decreased by 0, 1, or 2 (1/3 chance each). If they rejected the offer, the next offer increased by 0, 1, or 2 (1/3 chance for each option). Such contingency did not exist in the No Control condition, where the offers were randomly drawn from a Gaussian distribution (μ = 5, σ = 1.2, rounded to the nearest integer, max = 8, min = 2) and participants’ behaviors had no influence on future offers.

Against each team, participants played as the responder in 40 rounds of a social exchange game adapting the rule from the ultimatum game (Camerer, 2011). In the No Control condition, on each round, participants were offered a split of $20 from their partners (unbeknownst to participants, the actual offer was randomly drawn from a normal distribution, except the first offer which was always $5) and decided whether to accept or reject the offer. Here, participants’ current choices had no influence on the next offers from their partners. Critically, under the In Control condition, participants could exert control over their partners using their own actions. Specifically, participants’ current decisions (i.e. to accept or reject the offer) influenced the next offers from their partners in a systematic manner. Subject only to being between $1 and $9 (inclusive), partners increased the next offer by $0, $1, or $2 (probability of ⅓ each, subject to the constraints) if the participant rejected the present offer, and decreased the next offers by $0, $1, or $2 (probability of ⅓ each, again subject to the constraints) if the participant accepted the current offer (**Figure 1b** and **Methods**). Again, the starting offer was $5. On 60% of the trials, participants were asked about their emotional state after they made a choice (i.e., 24 ratings per condition; see **Figure S2**), and at the end of the task, they were asked to rate how much control they believed they had over their partners in each condition (using a 0-100 scale).

Note that participants were not instructed about the statistics of the task environment nor the nature of the condition they were playing; although the instruction about the existence of two separate teams was provided to encourage participants to learn contingent rules and norms within each condition. If participants were able to detect social controllability correctly within each condition, they would be expected to show strategic decisions that exert appropriate level of control over others’ subsequent choices.

## RESULTS

### Participants exerted control over others in controllable interactions

We first examined whether participants were able to detect the difference in controllability between the two social environments without explicit instruction. Our primary measures here were the offer sizes participants received in each condition; their rejection behaviors; and their self-reported level of perceived control. If individuals learned the contingency of the controllable condition, we should observe that 1) offers received under In Control would be pushed up to a higher level than those under No Control; 2) people would need to reject more offers to obtain larger future offers under In Control than No Control; and 3) people would report higher perceived control for In Control than for No Control.

First, we found that despite the same starting offer of $5, participants received higher offers over time under In Control compared to No Control condition (**Figure 2a1, 2**), indicating that individuals in general successfully exerted control over their partners without being explicitly instructed whether or not they had control.

**Figure 2.**
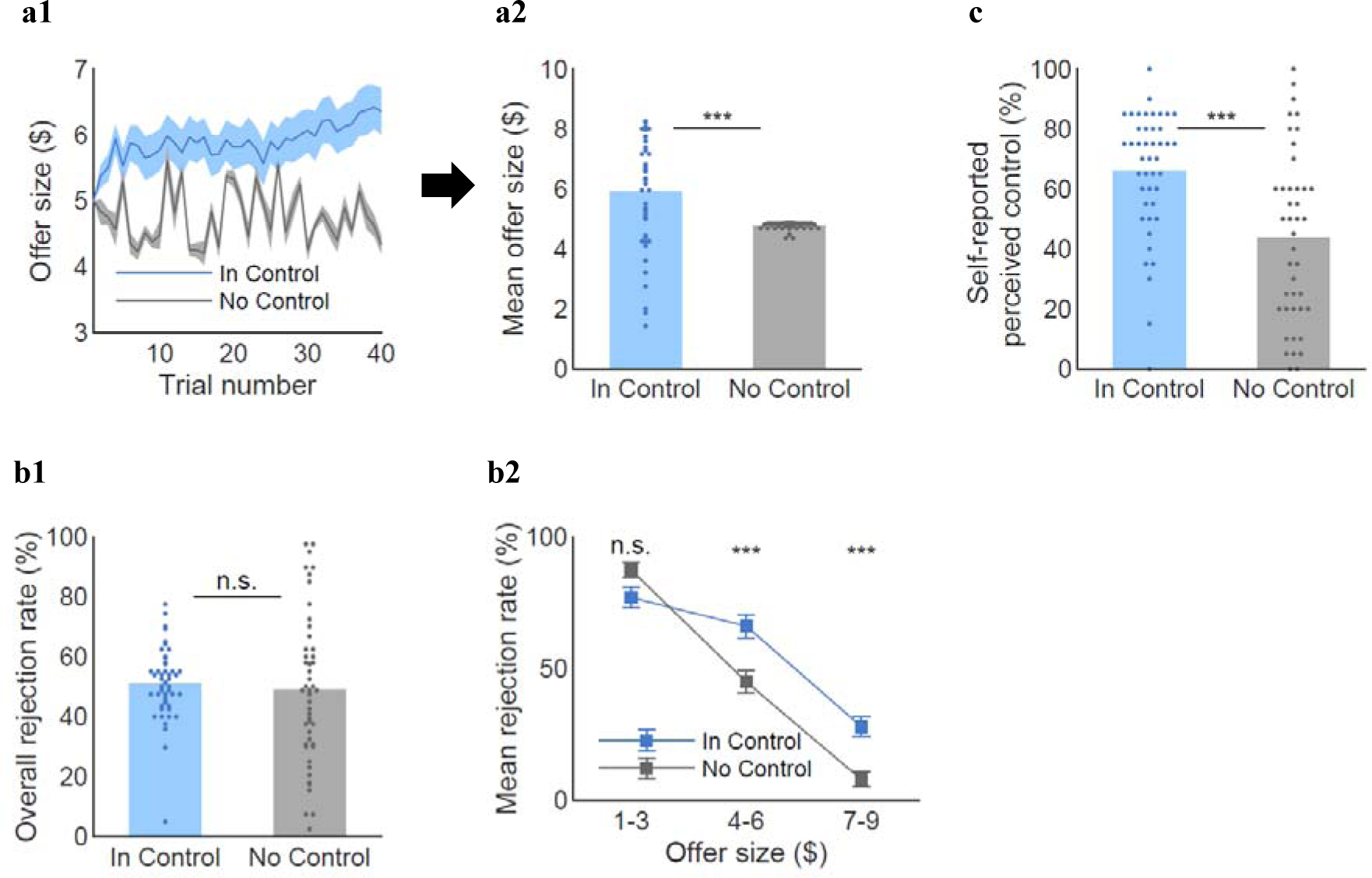
Model agnostic behavioral results. (**a1**) Participants raised the offers along the trials when they had control, compared to when they had no control. (**a2**) The mean offer size was higher for the In Control condition than the No Control condition (mean_ic_ = 5.9, mean_nc_ = 4.8, *t*(47.48) = 4.28, *P* < 0.001). (**b1**) The overall rejection rates were not different between the two conditions (mean_ic_ = 50.2%, mean_nc_ = 48.6%, *t*(67.75) = 0.39, *P* = 0.70). (**b2**) However, participants were more likely to reject middle ($4-6) and high ($7-9) offers when they had control (paired *t* test: *low($1-3)* mean_ic_ = 77%, mean_nc_ = 87%, *t*(22) = −1.38, *P* = 0.18, *middle($4-6)* mean_ic_ = 65%, mean_nc_ = 44%, *t*(47) = 5.26, *P* < 0.001, *high($7-9)* mean_ic_ = 28%, mean_nc_ = 8%, *t*(40) = 4.57, *P* < 0.001). Each offer bin for In Control condition in **c2** represents 23, 48, and 41 participants who were proposed corresponding offers at least once, whereas each bin for No Control condition represents all 48 participants. (**c**) The perceived control ratings were higher for In Control than No Control (mean_ic_ = 65.9, mean_nc_ = 42.3, *t*(76.99) = 4.48, *P* < 0.001; Six participants did not report perceived control ratings for at least one condition and were excluded from the ratings analysis). The t-statistics for the mean offer size, overall rejection rate, and perceived control are from two-sample *t* tests assuming unequal variance according to the results of the F-test for equal variance (offer: std_ic_ = 1.8, std_nc_ = 0.1, *F*(47, 47) = 194.38, *P* < 10^-40^; overall rejection rate: std_ic_ = 12%, std_nc_ = 25%, *F*(47, 47) = 0.23, *P* < 10^-5^; perceived control: std_ic_ = 20.3, std_nc_ = 28.8, *F*(45, 43) = 0.50, *P* < 0.05). Satterthwaite’s approximation was used for the effective degrees of freedom for t-test with unequal variance. The variance did not significantly differ for the binned rejection rates. Errorbars represent s.e.m.. \*\*\**P* < 0.001, n.s. indicates not significant.

Next, we examined the rejection patterns for both conditions. On average, rejection rates in the two conditions were comparable (**Figure 2b1**). By dissociating all trials in three levels of offer sizes (low: $1-3, medium: $4-6, and high: $7-9), we found that participants were more likely to reject medium to high ($4-9) offers when they had control, while they showed comparable rejection rates for the low offers ($1-3) between the two conditions (**Figure 2b2**). These results suggest that participants behaved in a strategic way to utilize their influence over the partners. One possible confound is that individuals may have experienced different affective states in the two conditions, thus changing choice behaviors. However, this seemed unlikely because there was no significant difference in emotional rating between In Control and No Control conditions (**Figure S2**).

As additional evidence for participants’ knowing and purposeful rejection behaved under In Control, we compared the self-reported belief about control between the two conditions. As expected, participants reported higher perceived control for In Control than No Control (**Figure 2c**), indicating that participants were aware of the difference in controllability between the two conditions. Nevertheless, the mean level of perceived control for No Control was 42.3%, which was still substantially higher than 0%. This result suggests that participants suffered an illusion of control when they had no control over their partners’ offers. Taken together, these findings demonstrate that participants learned controllability on their own and that they were able to raise the offers of the computer program by rejecting when they were given control, despite having developed some level of illusion of control. We investigate the mechanisms underlying these behaviors in the next sections.

### Participants used forward thinking to exert social control regardless of the actual controllability of the environment

Next, we sought to probe what cognitive processes would underlie such perception and behavior, by building computational models of participants’ choices. Previous studies on value-based decision-making have shown that people can use future-oriented thinking and mentally simulate future scenarios when their current actions have an impact on the future (Daw et al., 2011; Gläscher et al., 2010; Lee et al., 2014). Relying on this framework, we hypothesized that individuals use forward thinking to estimate their control over others and the values of exerting social control before making choices. To test this hypothesis, we constructed a set of forward thinking (FT) models which assume that an agent computes the values of action (here, accepting or rejecting) by summing up the current and the future values based on her estimation of the amount of control she has over the social interactions. These models also incorporate norm adaptation (Gu et al., 2015) to characterize how people calculate subjective values in social settings (Fehr and Schmidt, 1999) (versus non-social decision-making) and how to use these value signals to guide social choices (see **Methods** for details). The key individual-level parameter-of-interest in this model is estimated control, δ, representing the influence that participants thought they would have on the offer changes (see **Methods**). Moreover, we considered the number of steps one calculates into the future (i.e. planning horizon; **Figure 3a**). We compared models that considered one to four steps further in the future in addition to model-free reinforcement learning and standalone social learning both considering only the current step without forward thinking. In model fitting, we excluded the first five trials since we focused on the mechanism of *exerting* control rather than that of *learning* control. We also excluded the last five trials because the behavior could be different at the end of the episodes (e.g., “It is not worth trying to influence the partner on the 40th trial, since there are no future interactions”).

**Figure 3.**
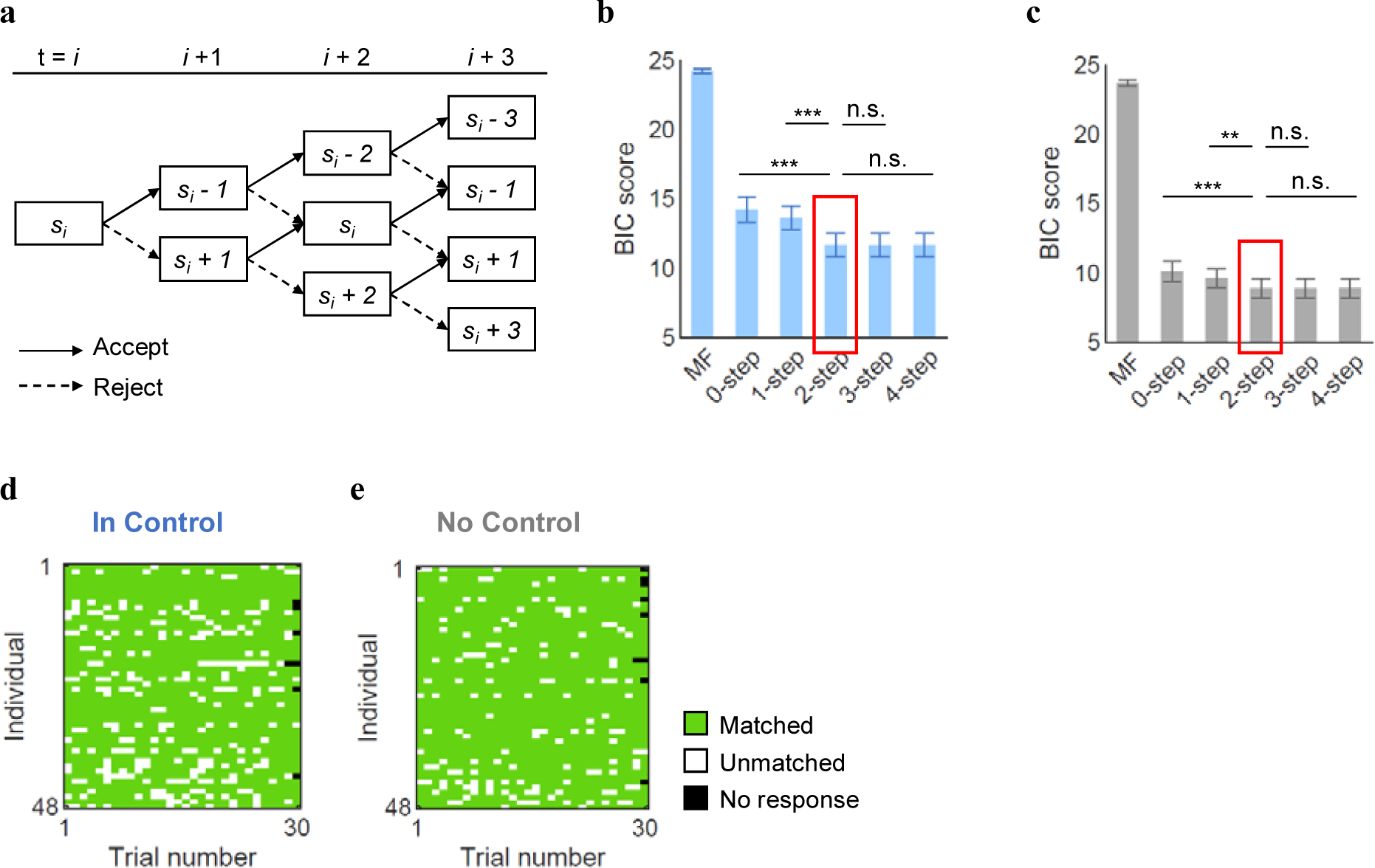
Computational modeling of social controllability. (**a**) The figure displays how the expected value of the offers evolve contingent on the choice along the future steps for In Control condition. To examine how many steps of horizon participants might simulate to exert control, we tested the candidate models considering zero to four steps of future horizon in addition to model-free learning. (**b-c**) For both In Control (**b**) and No Control (**c**) condition, the forward thinking (FT) models better explained participants’ behavior than the model-free learning and the no simulation model. The 2-step FT model was selected for the further analysis, as the marginal improvement in the BIC score drops for the longer simulation models (paired *t*-test comparing 2-step FT model with: (*i*) model-free learning IC *t*(47) = - 13.50, *P* < 10^-17^ NC *t*(47) = −18.61, *P* < 10^-22^; (*ii*) 0-step IC *t*(47) = −4.45, *P* < 0.0001 NC *t*(47) = −4.21, *P*< 0.001; (*iii*) 1-step IC *t*(47) = −4.41, *P* < 0.0001 NC *t*(47) = −3.01, *P* < 0.001; (*iv*) 3-step IC *t*(47) = 0.39, *P* = 0.70 NC *t*(47) = −0.04, *P* = 0.97; (*v*) 4-step IC *t*(47) = 0.06, *P* = 0.95 NC *t*(47) = −0.12, *P* = 0.91). The error bar represents the standard error of the mean. *** *P* < 0.001, ** *P* < 0.01. (**d-e**) The choices predicted by the 2-step FT model were matched with individuals’ actual choices with an average accuracy rate of (**d**) 83.7% for In Control (**e**) and 90.1% for No Control.

The results showed that for both conditions (In Control, No Control), all FT models significantly better explained participants’ choices than the model-free reinforcement learning and the standalone norm learning model without forward thinking (Gu et al., 2015), indexed by Bayesian Information Criteria (BIC) scores (**Figure 3b, c**). These results suggest that participants engaged in future-oriented thinking and specifically, calculated how their current choice might affect subsequent social interactions, regardless of the actual level of controllability of the environment. The FT models with longer planning horizon tend to show smaller BIC scores (i.e., better model fit), but the fit improvement became marginal after two steps (**Figure 4b, c**). The 2-step FT model predicted participants’ choices with an average accuracy rate of 83.7% for In Control (**Figure 4d**) and 90.1% for No Control (**Figure 4e**). Model recovery test also confirmed that all the parameters in the 2-step FT model were identifiable (**Figure S3a-j**). We thus used parameters from the 2-step FT model for subsequent neural analyses (see **Table 1** for a full list of parameters from this model).

**Table 1.** Parameter estimates from the 2-step forward thinking (FT) model.

|  | Inverse<br>temperature | Sensitivity<br>to norm<br>violation | Initial norm | Adaptation<br>rate | Estimated<br>control |
| --- | --- | --- | --- | --- | --- |
| | $\beta$ | $\alpha$ | $\mu$ | $\varepsilon$ | $\delta$ |
| **In Control** |  |  |  |  |  |
| Mean | 8.33 | 0.76 | 8.21 | 0.24 | 1.33 |
| SD | 8.55 | 0.29 | 7.14 | 0.24 | 0.79 |
| **No Control** |  |  |  |  |  |
| Mean | 10.38 | 0.79 | 8.84 | 0.29 | 0.98 |
| SD | 8.84 | 0.31 | 6.96 | 0.24 | 0.62 |

**Table 1. Parameter estimates from the 2-step forward thinking (FT) model.**
| | Inverse<br>temperature<br>$\beta$ | Sensitivity to<br>norm violation<br>$\alpha$ | Initial norm<br>$\mu$ | Adaptation<br>rate<br>$\varepsilon$ | Estimated<br>control<br>$\delta$ |
| --- | --- | --- | --- | --- | --- |
| **In Control** |  |  |  |  |  |
| Mean | 8.33 | 0.76 | 8.21 | 0.24 | 1.33 |
| SD | 8.55 | 0.29 | 7.14 | 0.24 | 0.79 |
| **No Control** |  |  |  |  |  |
| Mean | 10.38 | 0.79 | 8.84 | 0.29 | 0.98 |
| SD | 8.84 | 0.31 | 6.96 | 0.24 | 0.62 |

**Figure 4.**
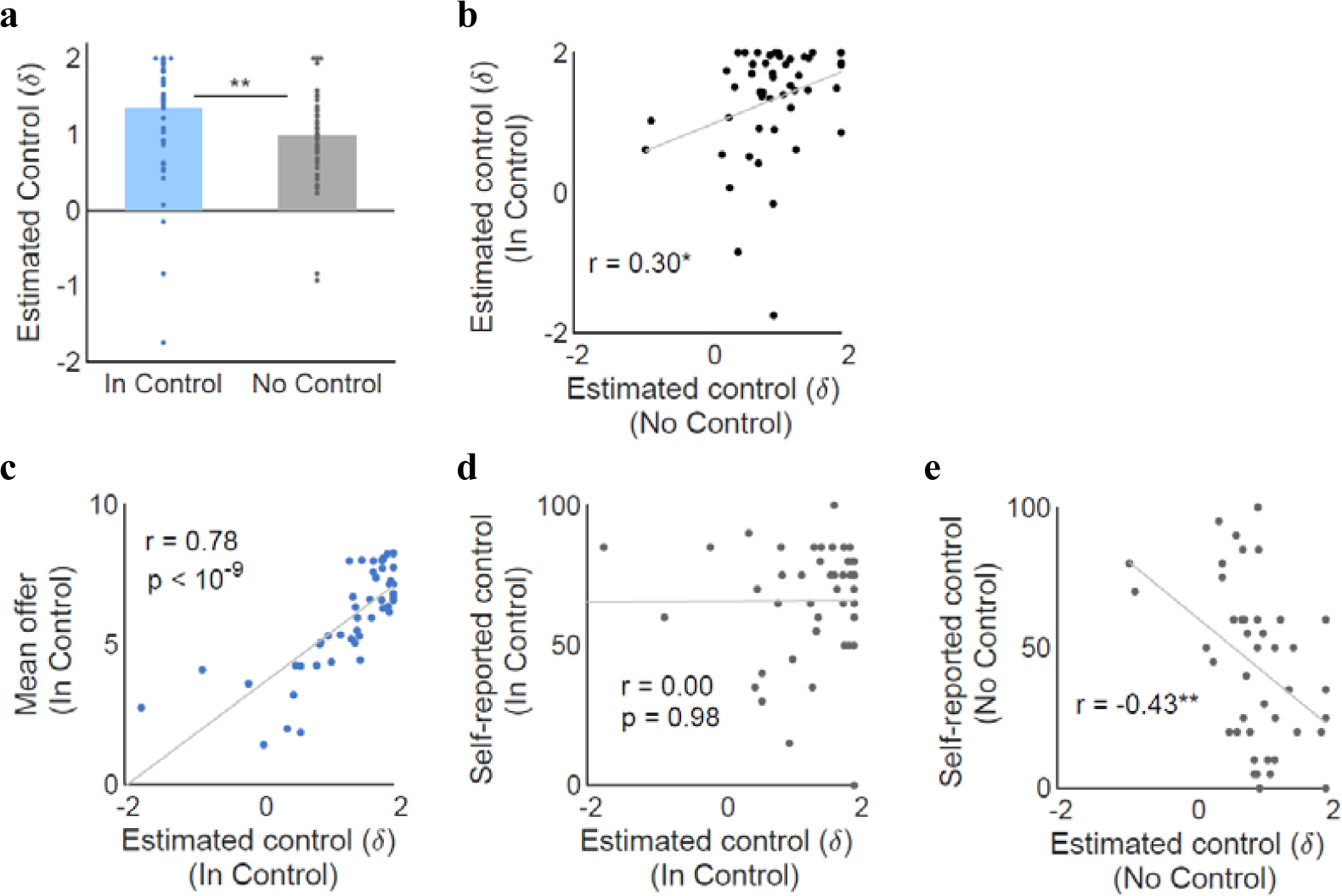
Estimated social control. (**a**) The levels of estimated control drawn from the 2-step forward thinking (FT) model were higher for In Control than for No Control (mean_ic_ = 1.33, mean_nc_ = 0.98, *t*(47) = 2.90, *P* < 0.01). (**b**) Estimated control showed trait-like characteristics in the sense that parameter estimates between the two conditions were positively correlated. (**c**) Under In Control, estimated control correlated with the mean offers. (**d**) Under In Control, estimated control was not significantly correlated with self-reported belief about control. (**e**) Under No Control, estimated control was anticorrelated with self-reported belief about control, suggesting that those who simulated higher level of control during the task ended up with more accurate perception of controllability. \**P* < 0.05, ** *P* < 0.01.

### The estimated control parameter δ captured trait-like characteristics

There are two possible interpretations about the estimated control δ, the key parameter in the 2-step FT model. First, the estimated control parameter δ could be a behavioral manifestation of illusion of control, consistent with the self-reported belief about control in the No Control environment shown in **Figure 2c**; or second, the estimated control δ could reflect one’s general tendency to have a sense of control. To examine these possibilities, we conducted four subsequent analyses regarding individually estimated δ from In Control and No Control conditions. First, we found that the estimated control estimates were higher for In Control than for No Control (**Figure 4a**), indicating that participants simulated greater levels of control when environments were in fact controllable than when they were uncontrollable. Second, δ estimates for In Control and No Control were correlated with each other (**Figure 4b**), indicating that the estimated control has trait-like characteristics. This was not the case for the self-reported belief about control (**Figure S4a**).

Third, under In Control, those who estimated greater control were more likely to raise the offers higher (**Figure 4c**), suggesting a positive association between estimated control and task performance during the In Control condition. In contrast, under the same In Control condition, there was no significant correlation between the estimated control and self-report belief about control (**Figure 4d**), or between self-reported beliefs and task performance (**Figure S4b**). Last, we found that under No Control, the estimated control parameter δ was anticorrelated with self-reported perceived control (**Figure 4e**). That is, those who estimated greater control during the task were less likely to end up with illusion of control at the end of the task. This result suggests that in uncontrollable social environments, using forward planning to estimate control helps detecting uncontrollability. While this finding seems counterintuitive at a first glance, we speculate that people with larger δ ended up experiencing larger prediction errors in the face of a lack of control, thus reporting reduced illusion of control at the end of the task. Taken together, this finding suggests that the δ parameter is *not* just another index of illusion of control and is unlike the self-reported beliefs; the estimated control parameter from the 2-step FT model reveals distinct individual characteristics that self-reports could not capture.

### Estimated values of social controllability were computed in overlapping, yet different neural regions in controllable and uncontrollable interactions

An important validation for any algorithm used by an information-processing system is the identification of the physical instantiation of the algorithm (e.g. the brain) (Marr and Poggio, 1976). Thus, to validate further that people did use forward thinking to exert social control, we examined whether decision values drawn from the FT model were computed in the brain. We regressed trial-by-trial simulated values of the chosen option drawn from the 2-step FT model as parametric modulators against event related blood-oxygen-level-dependent (BOLD) responses recorded during fMRI (see **Methods**). We found that the BOLD signals in the vmPFC tracked the value estimates in both In Control (**Figure 5a, Table S1)** and No Control conditions (**Figure 5b, Table S2**). This is consistent with the findings that vmPFC signals a common currency in value-based decision making in both social (Chung et al., 2015) and non-social domains (Levy and Glimcher, 2012). Additionally, the ventral striatum and anterior insula also encoded these value signals in the In Control condition (**Figure 5a, Table S1)**, but not in the No Control condition (**Figure 5b, Table S2)**. We further extracted the beta values from these ROIs for each condition (**Figure 5c-e**). This set of analyses suggested that indeed, vmPFC encoded value signals regardless of the actual controllability of the environment, and there was no difference in the beta estimates of the vmPFC between the two conditions (**Figure 5c**). In sharp contrast, both ventral striatum (**Figure 5d**) and anterior insula (**Figure 5e**) showed stronger encoding of these value signals in In Control, compared to No Control. Taken together, these neuroimaging results suggest that people used overlapping (i.e. vmPFC) yet different (e.g. striatum and insula) neural substrates to encode simulated social values in controllable and uncontrollable interactions.

**Figure 5.**
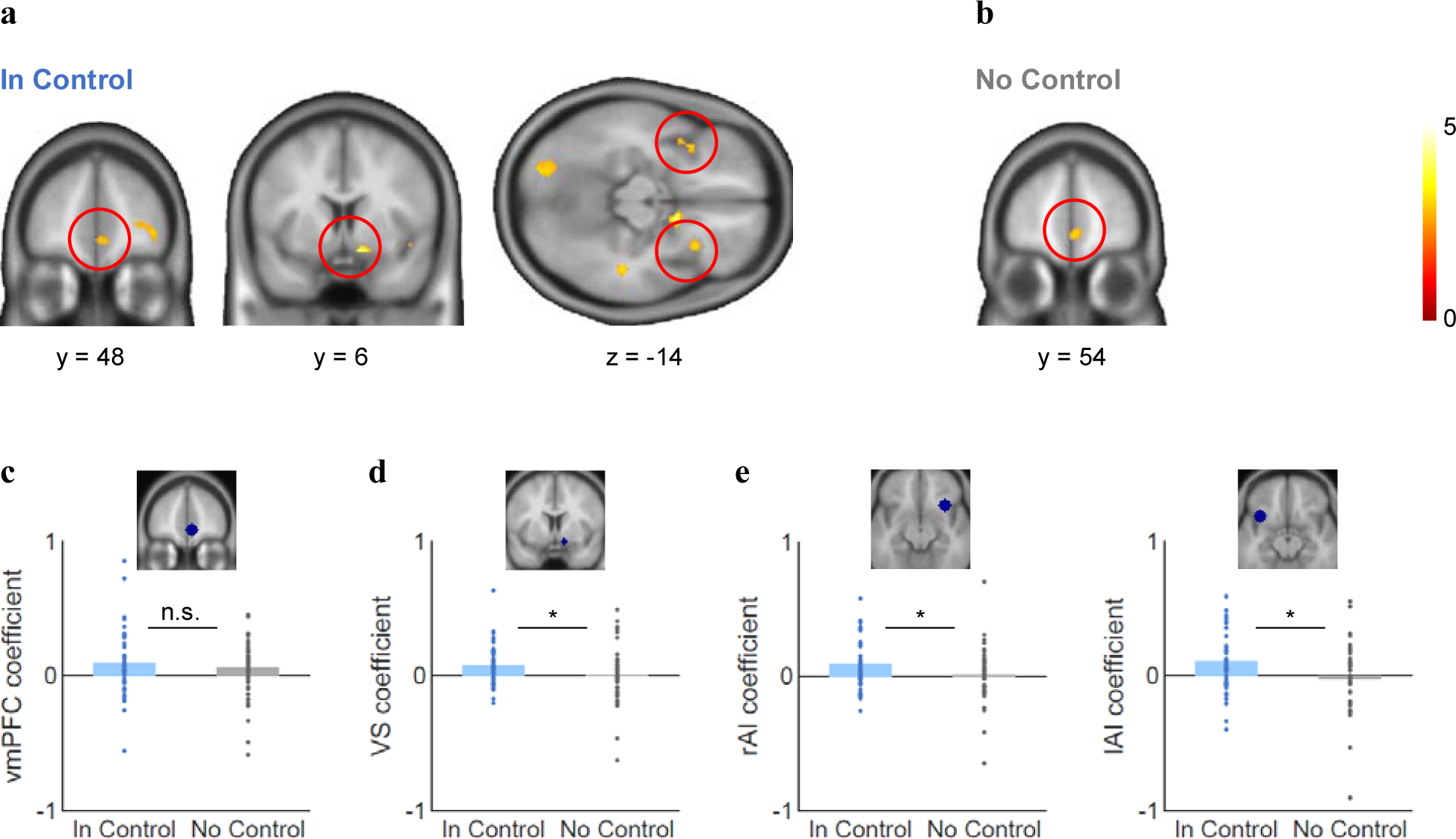
Neural computation of estimated values of social controllability. (**a**) For In Control (IC) condition, the ventromedial prefrontal cortex (vmPFC), ventral striatum, and insula encoded the mentally simulated values of the chosen actions drawn from the 2-step forward thinking (FT) model. (**b**) For No Control (NC) condition, only vmPFC, but not ventral striatum or insula encoded the estimated total values of the chosen actions drawn from the 2-step FT model. (Displayed at P < 0.005, uncorrected, k > 15. Cluster significance determined at P < 0.05 family-wise error (FWE), small volume correction). (**c**) ROI analyses showed that vmPFC encoding of simulated values was not statistically different between IC and NC (parameter estimate mean_ic_ = 0.09, mean_nc_ = 0.06, t(47) = 0.79, P = 0.43). **d-e**) ROI analyses also suggests that **d)** ventral striatum (parameter estimate mean_ic_ =, mean_nc_ =, t(47) = 2.09, P < 0.05,) and **e**) anterior insula value signals (parameter estimate right anterior insula mean_ic_ = 0.10, mean_nc_ = 0.01, t(47) = 2.26, P < 0.05; left anterior insula mean_ic_ = 0.11, mean_nc_ = −0.02, t(47) = 2.32, P < 0.05) were significantly greater for In Control than No Control.

## DISCUSSION

### Forward thinking as a computational mechanism for exerting social control

For social animals like humans, it is crucial to be able to identify situations where one is in control versus situations where the rules regulating strategic interactions are uncontrollable, and then to exert that social control when environment allows. The current study provides a mechanistic account for how people exert control over others during a social interaction. We demonstrate that 1) people are able to exert social control successfully through forward thinking, a mechanism captured by our computational model; 2) greater levels of control estimated by our participants correlated with better performance in controllable interactions and less illusion of control (e.g. self-reported beliefs) in uncontrollable interactions; and 3) values of social control were computed in the vmPFC, striatum, and insula for controllable interactions, but only in vmPFC for uncontrollable interactions. These findings demonstrate that people use forward thinking to guide social choices, a process implemented in various neural regions important for value-based decision-making.

Forward thinking is an important high-level cognitive process that is frequently associated with abstract reasoning (Hegarty, 2004), planning (Szpunar et al., 2014), and MB control (Constantinescu et al., 2016; Daw et al., 2011; Gläscher et al., 2010; Schuck et al., 2016; Wang et al., 2018). Also known as prospection, forward thinking has been suggested to involve four modes: mental simulation, prediction, intention, and planning (Szpunar et al., 2014). All four modes are likely to have taken place in our study, as our FT model implies that a social decision-maker mentally simulates social value functions into the future, predicts how her action would affect the following offers from partners, sets a goal of increasing future offers, and plans steps ahead to achieve the goal. Future studies will be needed to disentangle the neurocomputational mechanisms underlying each of these modes.

### Relevance to and advances over the MB and vmPFC literature

Critically relevant to the current study, previous research on MB control suggests that when faced with relatively novel, dynamically changing environments, decision-makers build and use an internal model or a cognitive map of simulated future states (Constantinescu et al., 2016; Daw et al., 2011; Gläscher et al., 2010; Schuck et al., 2016; Wang et al., 2018). The current study supports this MB control framework and further expands it to the social decision-making domain. Specifically, our 2-step FT model assumes that individuals not only use the current value, but also take the future consequences of their current actions into account to make social choices. Our model also has important advances over previous MB models, as it estimates how many steps people think forward and how much social control they have over partners, while incorporating aversion to norm violation and norm adaptation, two important parameters guiding social adaptation (Fehr, 2004; Gu et al., 2015; Spitzer et al., 2007). These individual and ‘social-specific’ parameters will be crucial for examining social deficits in various clinical populations in future studies.

Our modeling result was corroborated by neural findings of simulated social value encoding in the vmPFC across both conditions. In addition to its role in encoding ‘the common currency’ of values in non-social decision-making (Levy and Glimcher, 2012), the vmPFC has been shown to encode social decisions as well, such as computing ‘other-conferred utility’(Chung et al., 2015) and combining social values with reward-based learning signals (Behrens et al., 2008). More recently, this region has also been shown to represent mental maps of state space (Schuck et al., 2016) and of conceptual knowledge (Constantinescu et al., 2016), in addition to other ‘map’-encoding brain structures such as the hippocampus (O’keefe and Nadel, 1978; Tavares et al., 2015) and entorhinal cortex (Stensola et al., 2012). Thus, our results are consistent with these previous findings - and further expand the role of vmPFC to compute mentally estimated values of social control.

### Why do humans try to exert control in uncontrollable social situations?

One surprising finding from the current study is that people also used the 2-step FT model for the No Control condition, suggesting that they still estimated some level of control over their partners even when environment was in fact uncontrollable. At a first glance, estimated control derived from our model may seem as another index of subjective illusion of control (i.e. faulty belief that one has control even in uncontrollable environments). However, a closer examination suggests that the estimated control here reflects a dimension that is different from self-reported beliefs about control. Unlike the inconsistency of self-report beliefs (i.e. low correlation between belief ratings from In Control and No Control), estimated control (δ) from two different conditions were consistent within-subject, suggesting that this estimated control is capturing a trait-like characteristic. Furthermore, higher levels of estimated control in the No Control condition actually relates to reduced illusion of control, suggesting some potential positive outcomes of trying to exert control even in the No Control condition. We speculate that 1) people still attempted to simulate future interactions in uncontrollable situations due to their innate preference and tendency to control (Leotti and Delgado, 2014; Shenhav et al., 2016), and 2) those with higher levels of estimated control potentially experienced larger prediction errors and thus ended up with less illusion of control. Lastly, we also found that individuals who showed higher δ performed better in achieving higher offers from their partners under In Control, suggesting a direct association between forward thinking and performance in strategic social interaction. These results demonstrate that the forward thinking model and importantly the estimated control parameter captured a cognitive dimension that is distinct from self-reported beliefs about control, consistent with previous reports about the divergence between subjective and objective measures (e.g. (Sallis and Saelens, 2000; Shiffman et al., 2015)).

### Future directions

Given our results, it is compelling to design tasks that focus on the way that subjects learn the model (in our terms, acquiring a value for the parameter *δ*) in early trials or build complex models of their partners’ minds (as in a cognitive hierarchy (Camerer et al., 2004)). Indeed, even though, in our task, the straightforward model based on norm-adjustment characterized participants’ behavior well, there are more sophisticated alternatives that are used to characterize interpersonal interactions, such as the framework of interactive partially-observable Markov decision processes (Gmytrasiewicz and Doshi, 2005; Hula et al., 2015; Xiang et al., 2012). These might provide additional insights into the sorts of probing that our subjects presumably attempted in early trials to gauge controllability (and the ways this differs in both In and No Control conditions between subjects who do and do not suffer from substantial illusions of control). They would also allow us to examine whether our subjects thought that their partners built a model of them themselves (as in theory of mind or a cognitive hierarchy(Camerer et al., 2004)), which would add extra richness to the interaction, and allow us to capture individual trajectories more finely – if, for instance, our subjects might have become irritated (Hula et al., 2015) at their partners’ unwillingness to respond to their social signaling under No Control condition.

### Summary

The current study provides a mechanistic account for how people exert control over social others and how belief plays a role during social exchange. The implications of these findings could be far-reaching and multifaceted. First, the finding of engaging forward thinking in environments can help optimize normative social behavior, as often required during strategic social interaction (i.e. bargaining, negotiation). Second, our findings related to beliefs suggest that when beliefs are not grounded in reality, they can often be counter-productive in normative behavior, which are also a hallmark of many psychiatric disorders (e.g. schizophrenia). These findings might additionally imply that cognitive treatments that specifically target distorted beliefs may now be implemented and evaluated in a quantitative way.

## Acknowledgment

XG is supported by National Institute of Health grants R01DA043695, R21DA049243, R21 MH120789, and the Mental Illness Research, Education, and Clinical Center (MIRECC VISN 2) at the James J. Peter Veterans Affairs Medical Center, Bronx, NY. This study was supported by a faculty startup grant to XG from University of Texas, Dallas (where XG previously worked). DC is supported by UNIST internal funding (1.180073.01, 1.180031.01). PD is supported by the Max Planck Society.

## Author Contributions

Conceptualization: P.D. and X.G.; Methodology: S.N., D.C., A.H., P.D., and X.G.; Investigation: S.N. and J.J.; Formal Analysis: S.N. and D.C.; Writing – Original Draft: S.N. and X.G.; Writing – Review and Editing: S.N., D.C., A.H., V.F.G., P.D., and X.G.; Funding Acquisition: X.G.; Supervision: X.G.

## Declaration of Interests

The authors declare no competing interests.

## METHODS

### Participants

The study was approved by the Institutional Review Board of the University of Texas at Dallas and the University of the Texas Southwestern Medical Center (S.N., V.G.F, and X.G.’s previous institute where data were collected). Participants were recruited in the Dallas-Fort Worth metropolitan area. 56 healthy adults (38 female, age = 27.3 ± 9.2, 3 left-handed) provided written informed consent and completed this study. Five participants were excluded due to behavior data loss caused by computer collapse, one participant was excluded due to fMRI data loss, one participant was excluded due to excessive in-scanner head motion, and one participant was excluded due to poor quality of parameter recovery. The final sample had 48 healthy adults (33 female, age = 27.6 ± 9.1, 3 left-handed). Participants were paid a reward randomly drawn from the outcomes of this task, in addition to their baseline compensation calculated by time and travel distance.

### Task

We designed an economic exchange task to probe social controllability based on an ultimatum game. This task consisted of two blocks, each representing an experimental condition (‘In Control’ vs. ‘No Control’). In both conditions, participants were offered a split of $20 by a partner and decided whether to accept or reject the proposed offer from the partner. If a participant accepted the proposal, the participant and the partner split the money as proposed. If a participant rejected the proposal, both the participant and the partner received nothing. At the beginning of each block, participants were instructed that they would play the games with members of Team A or Team B. This instruction allows participants to perceive players in each block as a group with a coherent norm, rather than random individuals. However, participants were not told how the players in each team would behave so that participants would need to learn the action-offer contingency. There were 40 trials in each block. In 60% of the trials, participants were also asked to rate their feelings after they made a choice.

In the No Control condition, participants played a typical ultimatum game: the offers were randomly drawn from a truncated Gaussian distribution (μ = $5, σ = $1.2, rounded to the nearest integer, max = $8, min = $2) and participants’ behaviors had no influence on the future offers. Importantly, in the In Control condition, participants could increase the next offer from the partner by rejecting the current offer, or decrease the next offer by accepting the present offer in a probabilistic fashion (⅓ chance of ±$2, ⅓ chance of ±$1, ⅓ chance of no change; the range of the offers for In Control was between $1 and $9 (inclusive) – the range was not matched for the two conditions unintendedly.) (**Figure 1b**). We designed this manipulation based on the finding that reputation plays a crucial role in social exchanges(Fehr, 2004; King-Casas et al., 2005; Knoch et al., 2009); thus, in a typical ultimatum game, accepting any offer (although considered perfectly rational by classic economic theories(Becker, 2013)) will develop a reputation of being “cheap” and eventually lead to reduced offers, while the rejection response can serve as negotiation power and will force the partner to increase offers. At the end of each condition, participants also rated how much control they perceived using a sliding bar (from 0-100%).

### Computational modeling

We hypothesized that people would estimate their social controllability by using the consequential future outcomes to compute action values. To test this hypothesis, we constructed a forward-planning value function with different lengths of time horizons: zero to four steps of forward planning whereby zero-step represents the no planning model. For comparison purpose, we also considered model-free valuation(Gläscher et al., 2010). First, we assumed that participants correctly understood the immediate rules of the task as follows:

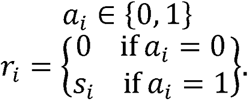

*a*_*i*_ represents the action that a participant takes at the *i*th trial where 0 representing rejection and 1 representing acceptance. *r*_*i*_ is the reward a participant receives at the *i*th trial depending on *a*_*i*_. Participants receive nothing if they reject whereas they receive the offered amount, *r*_*i*_, if they accept.

Similar to our previous work on norm adaptation(Gu et al., 2015), we assumed that people are averse to norm violations, defined as the difference between the actual offer received and one’s internal norm / expectation of the offers. Thus, the subjective utility of the expected immediate reward was constructed as follows.

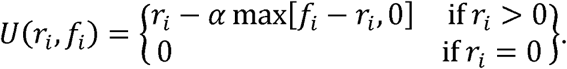

Here, *U*, the utility, is a function of the reward and *f* (“internal norm”) at the *i*th trial. The internal norm, which will be discussed in detail in the next paragraph, is an evolving reference value that determines the magnitude of subjective inequality. *α* (“sensitivity to norm violation”, 0 ≤ *α* ≤) 1represents the degree to which an individual is averse to norm violation. We assumed that if one rejected the offer and received nothing, aversion would not be involved as the individual already understood the task rule that rejection would lead to a zero outcome. Given that, if there is only one isolated trial, participants will choose to accept or reject by comparing *U*(*s*_*i*_, *f*_*i*_) and *U*(0, *f*_*i*_) = 0.

For the internal norm updating, as our previous study(Gu et al., 2015) showed that Rescorla-Wagner (RW)(Sutton and Barto, 2018) models fit better than Bayesian update models, we used RW norm updates to capture how people learn the group norm throughout the trials as follows.

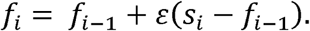

Here, *ε* is the norm adaptation rate (0 ≤ *ε* ≤), the individual learning parameter that determines the extent to which the newly observed offer is reflected to the posterior norm. The initial norm was set as a free parameter($ ≤ *f*_0_ ≤ 0$20).

Next, we formulated internal valuation as follows.

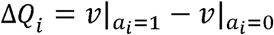

ΔQi, the difference between the value of accepting 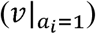 and the value of rejecting 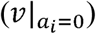, determines the probability of taking either action at the *i*th trial. Importantly, we incorporated forward thinking procedure in calculation of *v*. For a *n*-step forward thinking model, *v* was calculated as follows.

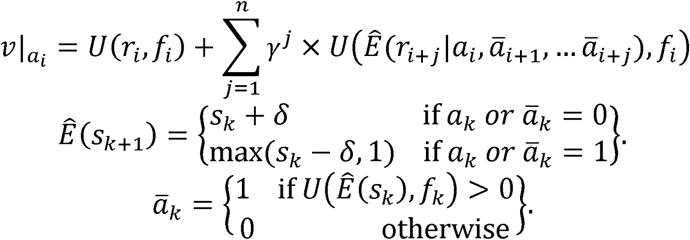

Given a hypothetical action *a*_*i*_ in the current (*i*^th^) trial, *v* is the sum of the expected future reward utility assuming simulated future actions, *ā*. We used the term *Ê* to represent an expected value in individuals’ perception and estimation. We assumed that in individual’s forward thinking, her hypothetical action at the future trial (*ā*_*k*_) increases or decreases the hypothetical next offer (*Ê*(*s*_*k*+1_) by *δ* (‘estimated control’, $2 ≤ *δ* ≤ $2). Here, we assumed symmetric change (*δ*) for either action so the change applies to both rejection and acceptance with the same magnitude but in the opposite direction. We assumed that simulated future actions (*ā*_*k*_) are deterministic contingent on the subjective utility of the immediately following rewards (*U*(*Ê*(*s*_*k*_),*f*_*k*_); this is a form of 1-level reasoning in a cognitive hierarchy(Camerer et al., 2004). The future values computed through estimated control was discounted by *γ*, the temporal discounting factor. We fixed *γ* at 0.8, the empirical mean across the participants from one initial around of estimation, in order to avoid collinearity with the parameter of our interest, *δ.*

We modeled the probability of accepting the offer using the softmax function as follows:

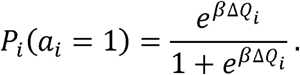

Here, *β* (‘inverse temperature’, 0 ≤ β ≤ 20) indicates how strictly people base their choices on the estimated value difference between accepting and rejecting. The lower the inverse temperature is, the more exploratory the choices are.

Model-free valuation was built as follows:

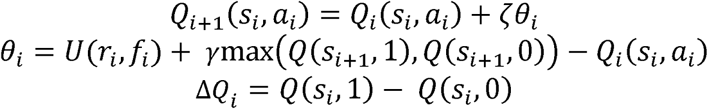

That is, after the reward (*r*_*i*_) and the next offer (*s*_*i*+1_) is observed, the value of the offer given the chosen action (*Q*(*s*_*i*_, *a*_*i*_) updates the reward prediction error (‘RPE’), *θ*_*i*_ with a learning rate of *ζ*(0 ≤ *ζ* ≤ 1). The actual rewards compared with the prediction in RPE consist of the utility of the immediate reward and the discounted value of the observed following offer, where the value of the following offer assumes deterministic greedy choice at the corresponding trial, consistent with the MB valuation. Δ*Q*_*i*_, the difference in values between accepting and rejecting, was entered to the softmax function in the same way as the MB valuation.

We fit the model to individual choice data for the middle 30 trials. We cut the first 5 trials in which one might be still learning the contingency between their action and the outcomes. Also, we excluded the last 5 trials since the room to increase the offers becomes smaller and participants had less incentive to reject offers, as the interactions were close to the end (Gneezy et al., 2003).

### fMRI data acquisition and pre-processing

Anatomical and functional images were collected on a Philips 3T MRI scanner. High-resolution structural images were acquired using the MP-RAGE sequence (voxel size =1 mm × 1 mm × 1 mm). Functional scans were acquired during the participants proceeded the task in the scanner. The detailed settings were as follows: repetition time (TR) = 2000 ms; echo time (TE) = 25 ms; flip angle = 90°; 38 slices; voxel size: 3.4 mm × 3.4 mm × 4.0 mm. The functional scans were preprocessed using standard statistical parametric mapping (SPM12, Wellcome Department of Imaging Neuroscience; www.fil.ion.ucl.ac.uk/spm/) algorithms, including slice timing correction, co-registration, normalization with resampled voxel size of 2mm × 2mm × 2mm, and smoothing with an 8mm Gaussian kernel. A temporal high-pass filter of 128 Hz was applied to the fMRI data and temporal autocorrelation was modeled using a first-order autoregressive function.

### fMRI general linear modeling

We specified GLM with a parametric modulator of the chosen actions’ values estimated from the 2-step FT model for each condition at the individual level using SPM12. The event regressors were 1) offer onset, 2) choice submission, 3) outcome onset, and 4) emotion rating submission. The parametric modulator was entered at the event of choice submission. In addition, six motion parameters were included as covariates. After individual model estimation, we generated the contrast images of whole-brain coefficient estimates of the chosen value. At the group level, we conducted a one-sample t-test for the simulated value regressor. Whole brain level analysis was thresholded at voxelwise *P* < 0.005 uncorrected and P < 0.05 family-wise error (FWE). Activations of regions of *a priori* interest (ROIs; including the vmPFC, striatum, and insula, based on their roles in value-based(Smittenaar et al., 2013) and social decision-making(Steinbeis et al., 2012)) were determined using small volume correction(Poldrack, 2007). For the ROI analysis for chosen value signals (**Figure 5c-e**), the ventral striatum (a 4mm-radius sphere centered at [12, 6, −14]), left anterior insula (a 8mm-radius sphere centered at [−36, 18, −14]), and right anterior insula (a 8mm-radius sphere centered at [32, 20, −12]) were taken from the peak voxels for chosen values under In Control condition. ROIs were extracted using the MarsBaR toolbox (Brett et al., 2002).

