## Supplementary Information for "Humans Use Forward Thinking to Exert Social Control"

**Supplemental Information**

**a2**

**a1**

**c2**

**b**

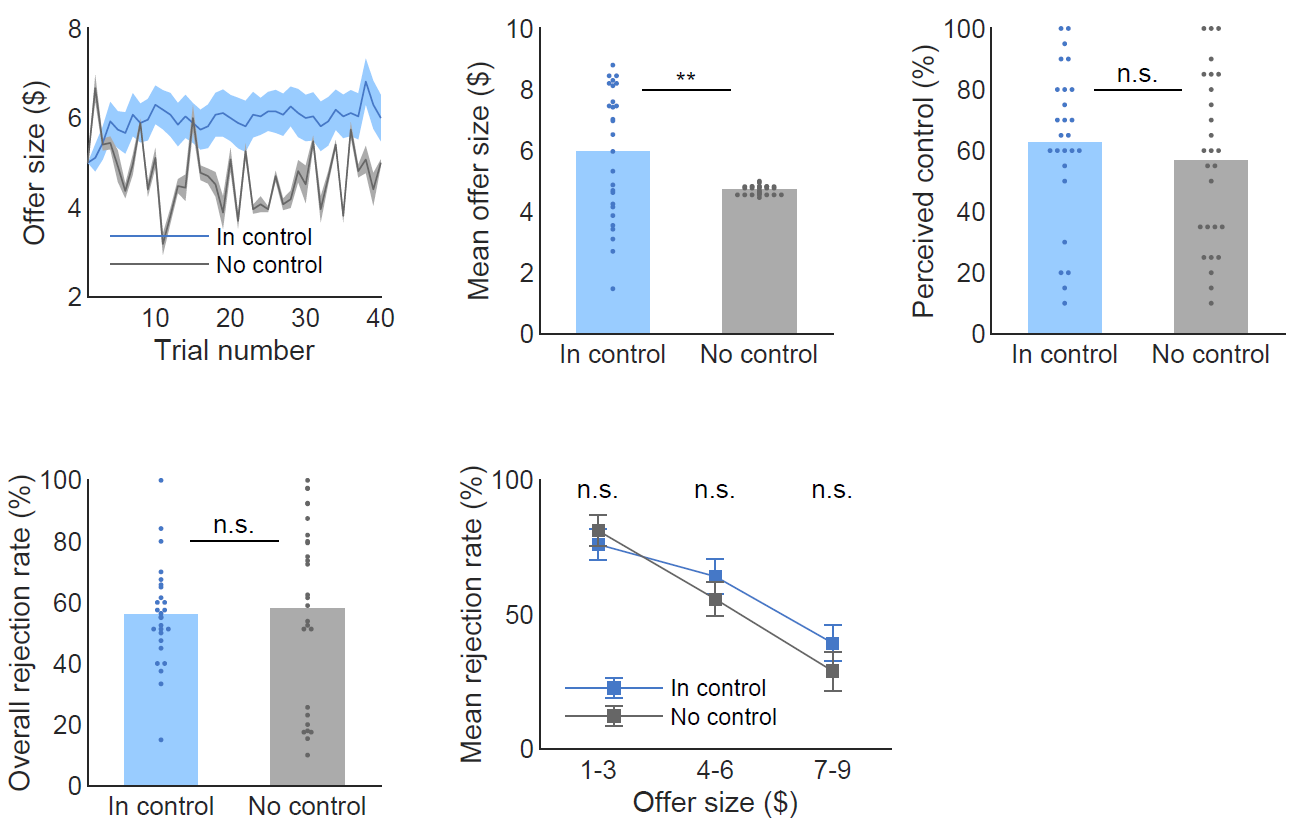

**c1**

**Figure S1. Behavioral results of the non-social task.** To investigate whether our results are specific to the social domain, we ran another batch of the task in which 27 out of the 48 original participants played the same game with the instruction of “playing with computer” instead of “playing with virtual human partners”. (**a1-2**) The offer (mean_ic_ = 6.0, mean_nc_ = 4.7, *t*(26.23) =3.03, *P* < 0.01) was higher for In Control than No Control, similar to the results of the social task. However, (**b**) the self-reported perception of control was not different between the two conditions when individuals played the game with a computer (mean_ic_ = 62.7, mean_nc_ = 56.9, *t*(25) =0.78, *P* = 0.44). (**c1**) The overall rejection rates (mean_ic_ = 55.9%, mean_nc_ = 58.1%, *t*(40.76) = -0.33, *P* = 0.74) or (**c2**) any of the binned rejection rates were not significantly different between the two conditions (paired *t* test: *low($1-3)* mean_ic_ = 76%, mean_nc_ = 81%, *t*(12) = 1.54, *P* = 0.15, *middle($4-6)* mean_ic_ = 64%, mean_nc_ = 56%, *t*(26) = 1.74, *P* = 0.09, *high($7-9)* mean_ic_ = 39%, mean_nc_ = 29%, *t*(19) = 0.80, *P* = 0.44). Satterthwaite’s approximation was used for the effective degrees of freedom for t-test with unequal variance. The variance significantly differed for the offer and the overall rejection rates. Errorbars represent s.e.m.. ** *P* < 0.01, n.s. indicates not significant.

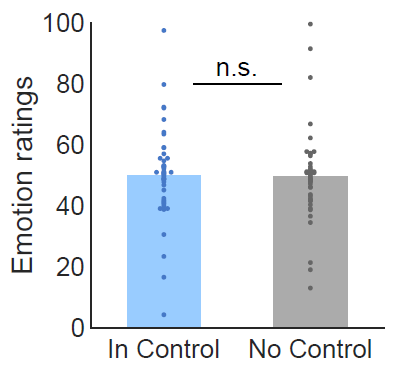
`

**Figure S2. Emotion ratings.** The mean emotion ratings were not significantly different between the two conditions (mean_ic_ = 49.9, mean_nc_ = 49.7, *t*(47) = 0.10, *P* = 0.92).

**a**

**b**

**c**

**d**

**e**

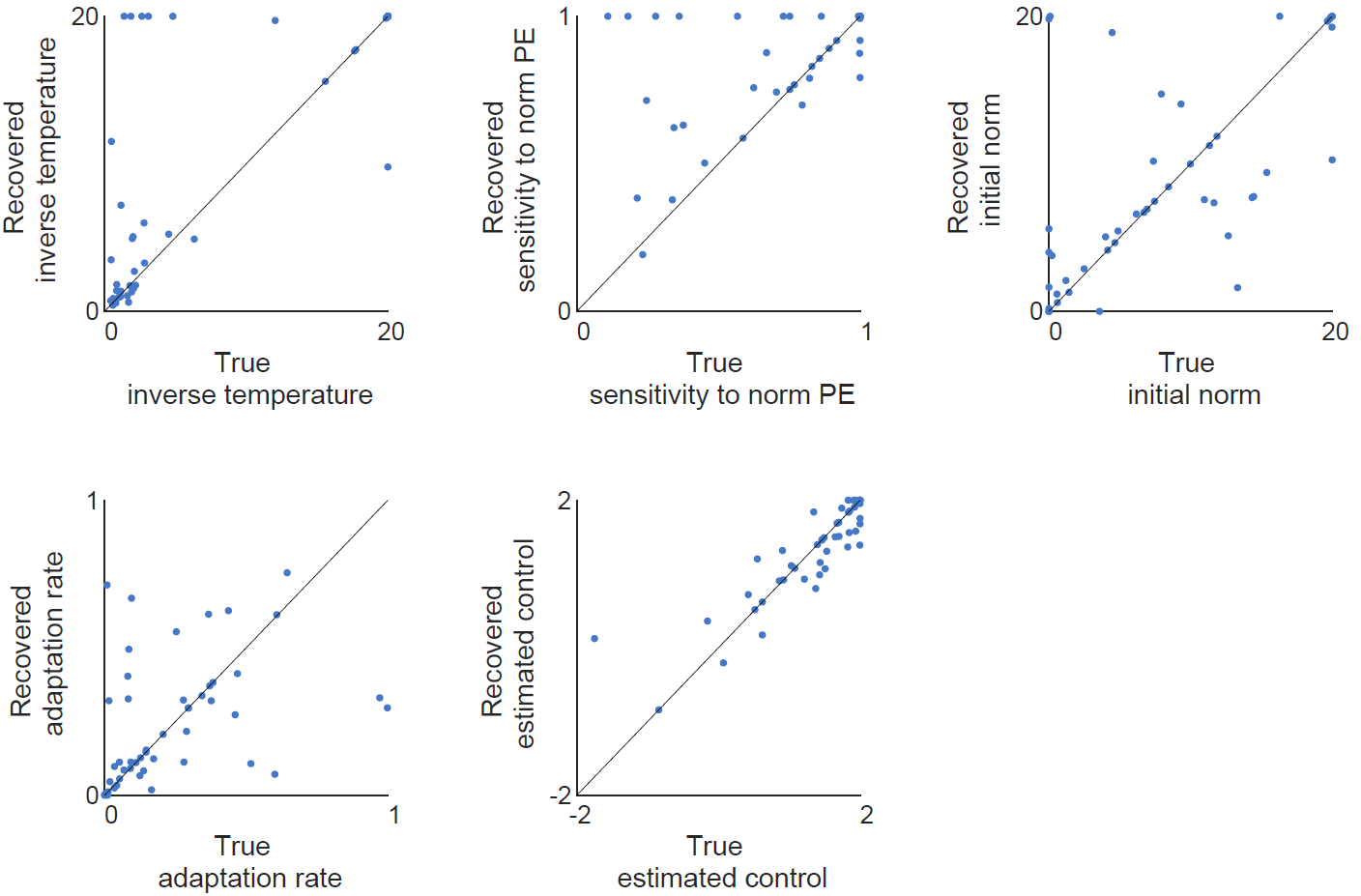

In Control

No Control

**f**

**g**

**h**

**i**

**j**

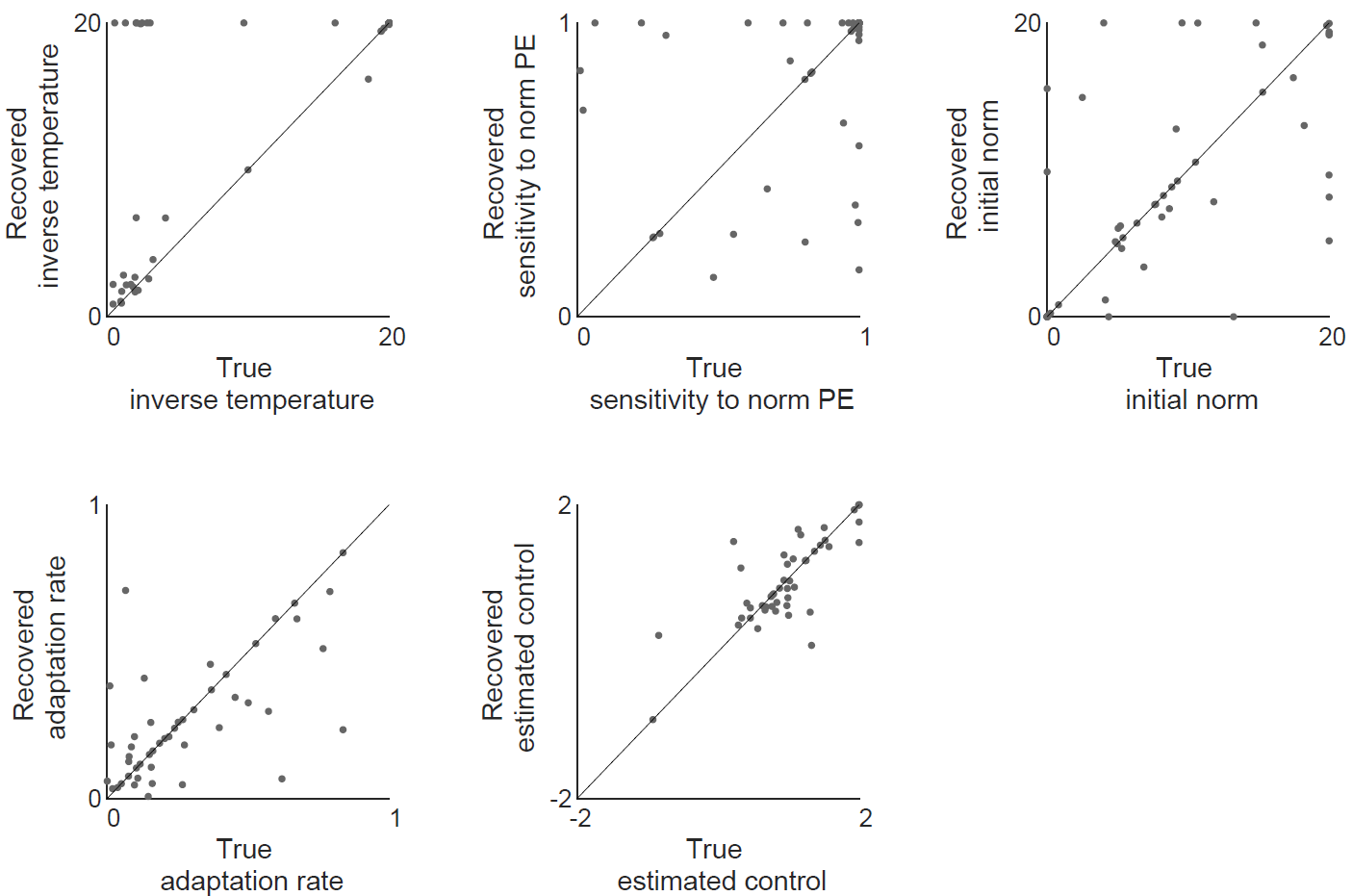

**Figure S3. Parameter recovery.** To assess the estimation quality of the 2-step FT model, we tested whether the parameters are precisely recovered for both conditions. We set the parameter estimates from the model as the true parameters and then we generated individual’s response data given the actual offers and those true parameters, using the same model. Under In Control (**a**-**e**), all parameters, (**a**) the inverse temperature (γ) (*R* = 0.77, *P*- < 10^-9^), (**b**) sensitivity to norm PE (α) (*R* = 0.57, *P* < 10^-4^), (**c**) initial norm (μ) (*R* = 0.66, *P* < 10^-6^), (**d**) adaptation rate (ε) (*R* = 0.39, *P* < 0.01), and (**e**) estimated control (δ) (*R* = 0.88, *P* < 10^-15^) were well identified as shown here. Under No Control as well, all parameters, (**f**) the inverse temperature (γ) (*R* = 0.69, *P*- < 10^-7^), (**g**) sensitivity to norm PE (α) (*R* = 0.33, *P* < 0.05), (**h**) initial norm (μ) (*R* = 0.62, *P* < 10^-5^), (**i**) adaptation rate (ε) (*R* = 0.66, *P* < 10^-6^), and (**j**) estimated control (δ) (*R* = 0.79, *P* < 10^-10^) were well identified. Each dot represents one individual.

| **a** | **b** |
| --- | --- |
| 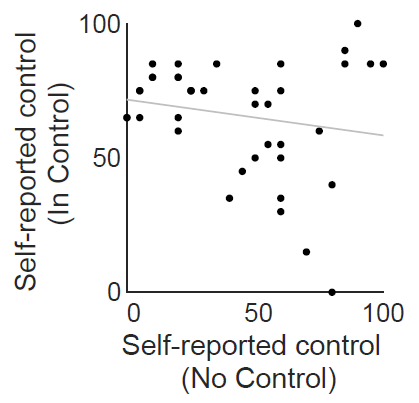 | 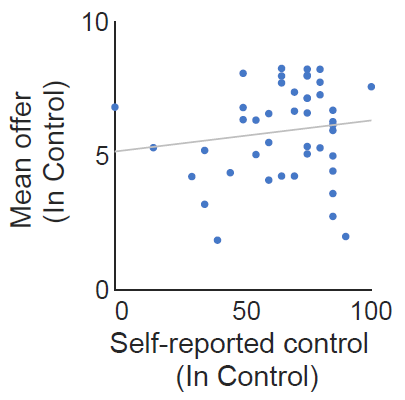 |

**Figure S4. Self-reported beliefs about control.** (**a**) Self-reported perceived control was not significantly correlated between In Control and No Control (*R* = -0.18, *P* = 0.26). (**b**) Under In Control, self-reported perceived control was not correlated with the mean offer size (*R* = 0.14, *P* = 0.37).

**Table S1. Brain activations related to chosen values (In Control)**

| Region | Lat | x | y | z | T | Z | k |
| --- | --- | --- | --- | --- | --- | --- | --- |
| Postcentral gyrus | R | 36 | -22 | 50 | 5.02 | 4.47 | 883 |
| Ventral striatum* | R | 12 | 6 | -14 | 4.38 | 3.99 | 25 |
| Inferior occipital gyrus | R | 20 | -88 | -8 | 3.93 | 3.63 | 17 |
| Cerebellum | L | -22 | -82 | -20 | 3.73 | 3.47 | 167 |
| Cerebellum | L | -10 | -62 | -50 | 3.62 | 3.38 | 43 |
| Cerebellum | L | -22 | -52 | -24 | 3.35 | 3.15 | 155 |
| Inferior frontal gyrus, pars orbitalis | R | 40 | 36 | -4 | 3.32 | 3.13 | 50 |
| Insula* | R | 32 | 20 | -12 | 3.25 | 3.07 | 124 |
| Inferior occipital gyrus | R | 42 | -80 | -8 | 3.03 | 2.88 | 18 |
| Middle frontal gyrus | R | 30 | 50 | 6 | 3.02 | 2.88 | 70 |
| Inferior temporal gyrus | R | 52 | -52 | -8 | 3 | 2.86 | 18 |
| Ventromedial prefrontal cortex* | R | 6 | 48 | -6 | 2.96 | 2.82 | 38 |
| Insula* | L | -36 | 18 | -14 | 2.92 | 2.79 | 19 |

(*P*<0.005 uncorrected and *K*>15; **P* < 0.05 corrected for small volume)

**Supplemental Table 2. Brain activations related to chosen values (No Control)**

| Region | Lat | x | y | z | T | Z | k |
| --- | --- | --- | --- | --- | --- | --- | --- |
| Inferior parietal lobule | R | 52 | -24 | 46 | 5.78 | 5 | 17938 |
| Cerebellum | L | -30 | -48 | -46 | 4.16 | 3.82 | 192 |
| Posterior cingulate gyrus | R | 2 | -44 | 28 | 3.43 | 3.23 | 381 |
| Inferior frontal gyrus, pars triangularis | R | 50 | 34 | 14 | 3.37 | 3.17 | 198 |
| Middle temporal gyrus | L | -54 | -34 | -12 | 3.31 | 3.12 | 23 |
| Inferior frontal gyrus, pars orbitalis | R | 56 | 4 | 12 | 3.28 | 3.1 | 30 |
| Precentral gyrus | L | -38 | 0 | 34 | 3.28 | 3.1 | 49 |
| Ventromedial prefrontal cortex* | R | 2 | 54 | -6 | 3.27 | 3.09 | 38 |
| Supplementary motor area | R | 2 | 8 | 54 | 3.2 | 3.02 | 56 |
| Precentral gyrus | R | 48 | 8 | 38 | 3.16 | 3 | 140 |
| Inferior frontal gyrus, pars triangularis | L | -48 | 28 | 32 | 3.08 | 2.92 | 15 |

(*P*<0.005 uncorrected and *K*>15; **P* < 0.05 corrected for small volume)

**Table S3. Contrast between the two conditions (IC > NC)**

| Region | Lat | x | y | z | T | Z | k |
| --- | --- | --- | --- | --- | --- | --- | --- |
| Temporal pole | L | -42 | 16 | -16 | 3.34 | 3.15 | 607 |
| Ventral Striatum | R | 10 | 6 | -14 | 3.11 | 2.95 | 729 |
| Inferior frontal gyrus, pars orbitalis | R | 48 | 38 | -8 | 2.8 | 2.68 | 1320 |
| Insula | R | 30 | 16 | -14 | 2.78 | 2.67 | - |

(*P* < 0.05 uncorrected, *K*> 120)

**Table S4. Contrast between the two conditions (NC > IC)**

| Region | Lat | x | y | z | T | Z | k |
| --- | --- | --- | --- | --- | --- | --- | --- |
| Middle temporal gyrus | L | -50 | -74 | 16 | 4.45 | 4.04 | 18606 |
| SupraMarginal gyrus | R | 48 | -36 | 26 | 3.35 | 3.16 | 1068 |
| dlPFC* | L | -36 | 0 | 32 | 2.94 | 2.81 | 574 |
| Middle frontal gyrus | L | -36 | 38 | 36 | 2.62 | 2.52 | - |
| Inferior frontal gyrus, pars triangularis | L | -44 | 20 | 24 | 2.56 | 2.47 | - |
| Precentral gyrus | R | 50 | 0 | 32 | 2.94 | 2.81 | 370 |

(*P* < 0.05 uncorrected, *K*> 120)

**Notes S1. Task instructions.** We provided participants written instructions as below.

Task Instructions (Compensation $0-10)

In this task you will participate in a series of decisions to split $20.00 with a partner. Your partner will propose how to split the money. **You will decide whether or not to accept the proposal.** If you accept the proposal, you will each get the share of the split your partner proposed. If you reject the proposal, however, both of you will get nothing.

At the beginning of each round you will be randomly paired up with a **new** partner. Because this is done randomly, you will be playing a different partner in each round. You will not meet any of your partners at any point throughout the session.

You will play total two rounds in a random order and each group has 40 players.

**Here is what the game will look like:**

You will be assigned a group of proposers to play the game with.

You will be playing with proposers from Group A

**HIT SPACE TO CONTINUE**

At the beginning of each round, you will be randomly paired with a partner and you will see their proposal to split money between you and your partner:

H.T. proposes you: $6

You can choose to accept the proposed split or reject the offer.

H.T.	YOU

$14	$6

<- ACCEPT	REJECT->

You receive $6

After some trials you will be asked how you feel about the proposal. Use left and right arrow keys to move the slider bar. To submit your choice, press the space bar.

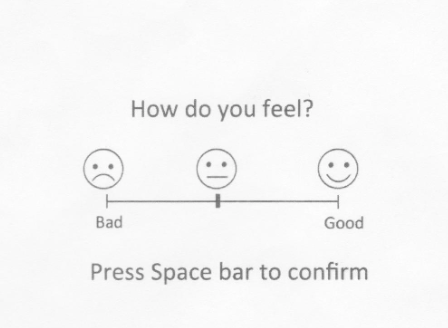

If you reject the offer, both you and your partner will not earn any money.

After you see the result of the proposal, you will be randomly paired with a new partner and a new round will begin. You will play **80** rounds of this game in total.

BONUS PAYMENT: At the end of the experiment the computer will randomly draw one of the rounds you played and pay you a bonus according to the outcome of that round: your share of the money if your partner accepted the offer, or nothing if they rejected the offer. Because any **one** of the choices could count “for real”, we encourage you to make each choice as though it is the one you are actually going to receive.
